# Acquired olfactory loss alters functional connectivity and morphology

**DOI:** 10.1101/2021.01.11.426175

**Authors:** Behzad Iravani, Moa G. Peter, Artin Arshamian, Mats J. Olsson, Thomas Hummel, Hagen H. Kitzler, Johan N. Lundström

## Abstract

Removing function from a developed and functional sensory system is known to alter both cerebral morphology and functional connections. To date, a majority of studies assessing sensory-dependent plasticity have focused on effects from either early onset or long-term sensory loss and little is known how the recent sensory loss affects the human brain. With the aim of determining how recent sensory loss affects cerebral morphology and functional connectivity, we assessed differences between individuals with acquired olfactory loss (duration 7-36 months, n=20) and matched healthy controls (n=23) in their grey matter volume, using multivariate pattern analyses, and functional connectivity, using dynamic connectivity analyses, within and from the olfactory cortex. Our results demonstrate that acquired olfactory loss alters grey matter volume in, among others, posterior piriform cortex, a core olfactory processing area, as well as the inferior frontal gyrus and angular gyrus. In addition, compared to controls, individuals with acquired anosmia displayed significantly stronger dynamic functional connectivity from the posterior piriform cortex to, among others, the angular gyrus, a known multisensory integration area. No significantly stronger connectivity in healthy control participants were demonstrated. When assessing differences in dynamic functional connectivity from the angular gyrus, individuals with acquired anosmia had stronger connectivity from the angular gyrus to areas primary responsible for basic visual and taste processing. These results demonstrate that recently acquired sensory loss alters both cerebral morphology within core olfactory areas and increase dynamic functional connectivity from olfactory cortex to cerebral areas processing multisensory integration.

## INTRODUCTION

Sensory loss is thought to alter both functional neural connectivity and morphology within primary sensory-related areas due to the resulting deprivation of input (Frasnelli et al., 2011). Most evidence of sensory loss-related neuroplasticity originates from the visual and auditory systems where significant effects are demonstrated in primary sensory processing areas (Fine and Park, 2018; Röder and Rösler, 2004). Studies assessing impact on primary olfactory cortex (piriform cortex) due to loss of the sense of smell (anosmia) have, in contrast, reported either minor (Frasnelli et al., 2013; Karstensen et al., 2018; Peng et al., 2013; Reichert and Schöpf, 2018) or indiscernible functional or morphological changes (Han et al., 2018, 2017; Peter et al., 2021, 2020; Yao et al., 2018). The vast majority of studies exploring neural impact of acquired visual and auditory sensory loss have focused on adult individuals that experienced their sensory loss early, often before their second year of life, and with many years without sensory function. In contrast, the majority of studies exploring impact from olfactory loss have studied adult individuals with lifelong sensory loss or individuals that, divergent to the sensory loss literature at large, lived very short time with their sensory loss, often within a year of sensory insult.

As stated above, the vast majority of studies exploring neural impact of acquired visual and auditory sensory loss have focused on adult individuals with either early or lifelong sensory dysfunction. These individuals demonstrate greater behavioral alterations as well as greater functional and structural cortical reorganization, as compared to individuals with a sensory loss acquired later in life (Gougoux et al., 2004; Voss, 2013; but see, Goldreich and Kanics, 2003). In essence, it has been suggested that cortical reorganization due to sensory loss is minor if the loss occurred after early childhood. This limited reorganization has been explained by the fact that late sensory loss generally occurs after the so called sensitive periods early in life, during which sensory experience has a strong influence on behavioral and cortical development (Singh et al., 2018; Voss, 2013). In contrast, sensory ablation studies performed in adult non-human animal models demonstrate that neural reorganization of sensory cortices can be detected after shorter spans of sensory deprivation, as early as 16 days past total sensory loss, even though the animals are clearly past the sensitive periods (Allman et al., 2009; Buonomano and Merzenich, 1998). Furthermore, artificially induced, very short sensory deprivation (e.g., by blindfolding, ranging from an hour to a few days) in adult humans has been demonstrated to affect both cortical excitability (Boroojerdi et al., 2000; Fierro et al., 2005; Leon-Sarmiento et al., 2005; Pitskel et al., 2007) and crossmodal processing (Fierro et al., 2005; Pascual-Leone and Hamilton, 2001). Hence, on the one hand, effects of long lasting late-onset sensory deprivation seem to be small in the visual and auditory systems; on the other hand, significant neural plasticity effects can be demonstrated already after a few hours of artificially induced sensory deprivation.

Anosmia is primarily a late-onset sensory loss, mainly affecting middle-aged and elderly populations (Boesveldt et al., 2017). Combined with the fact that anosmia is our most common sensory loss, affecting an estimated full 5% of the population (Brämerson et al., 2004; Landis et al., 2004), and with the COVID-19 pandemic and its associated olfactory problems potentially significantly increasing these numbers even more (Iravani et al., 2020; Parma et al., 2020), anosmia is a good model to study the effects of intermediate or late-onset sensory loss on neural plasticity. For morphology, there is clear evidence of structural reorganization following anosmia in large part around the orbito-frontal cortex (Bitter et al., 2010; Han et al., 2017; Li et al., 2020; Peter et al., 2020; Yao et al., 2018) but also in areas outside what is commonly regarded as olfactory processing areas, such as the insula cortex (Han et al., 2018, 2017; Peng et al., 2013; Yao et al., 2014). In contrast, few studies have explored effects of olfactory sensory loss on functional connectivity from the primary olfactory cortex, the piriform cortex, or functional processing within. Whereas lifelong absence of olfactory input, so-called congenital anosmia, does not seem to alter functional connectivity either within or from piriform cortex during rest (Peter et al., 2021), acquired anosmia has been linked to decreased connectivity within the piriform cortex odor- and sniff-induced network (Kollndorfer et al., 2015; Reichert et al., 2018). Moreover, artificially induced shortterm olfactory loss by naris occlusion alters odor-related processing in both the posterior piriform cortex and the orbitofrontal cortex (Wu et al., 2012); areas linked to the formation of odor representation as well as, among other skills, odor attention (Lundström et al., 2011). Taken together, although a minor number of publications exist in comparison, results from odor-based sensory loss studies indicate largely inconsistent results compared to those obtained in studies assessing plastic effects originating from visual deprivation.

The few studies exploring plastic changes due to acquired anosmia have demonstrated inconsistent changes in cerebral function or structure of the primary olfactory cortex, the piriform cortex. In respect of function, this can be explained by the dearth of studies assessing effects but also potentially by the reliance on task-induced measures for functional connectivity analyses where the latter has been dependent on either odor or active sniff activity. Reliance on these tasks brings with it inherent problems. If odors are presented during data acquisition, healthy individuals will differ from individuals with anosmia in respect of their cognitive odor associations and other odor evaluations that extend beyond basic sensory processing of the presented odor. If a sniff task is used, healthy individuals might differ from individuals with anosmia in their readiness to perceive odors and hence produce expectation, potential search behaviors, or trying to imagine the odors (Arshamian and Larsson, 2014; Zelano et al., 2005). On the other hand, functional MRI in the absence of a task, so-called restingstate scanning, shows activation patterns similar to those evident during a task and is correlated with underlying structural connectivity (Hermundstad et al., 2013; Honey et al., 2009). Assessing plastic changes due to olfactory sensory loss between individuals with anosmia and healthy individuals using resting-state connectivity in the absence of olfactory stimuli is therefore a method that does not potentially confound results due to the task at hand. It has been hypothesized that the lack of clear sensory loss-related effects in the olfactory system might originate from the fact that the olfactory system is dependent on areas that are more heterogeneous with less sensory specialization than, for example, the visual and auditory systems (Small, 2004). If this is the case, multivariate focused analyses methods assessing interconnected changes would be more appropriate to assess potential insults rather than classical analysis methods that assess each analysis unit independently. With a few notable exceptions (Chen et al., 2020; Kollndorfer et al., 2015), studies assessing plastic changes in morphology and functional connectivity due to olfactory sensory loss have, however, to a large degree relied on individual voxel-dependent changes in morphology and static functional connectivity, respectively, and not network-based analyses where change over multiple voxels or dynamic connectivity are assessed.

To determine whether olfactory loss within 3 years of the insult, in other words in the time range between immediate and long-term sensory loss, cause neural reorganization, we first assessed potential sensory deprivation-dependent changes in grey matter volume using a support vector machine with a searchlight procedure to generate a wholebrain map of classification accuracy of differentiating individuals with acquired anosmia from healthy control individuals. In the same individuals, we then assessed potential sensory deprivation-dependent changes in dynamic functional connectivity from olfactory related areas.

## METHOD

### Participants

A total of 43 participants were included whereof 20 participants were diagnosed with functional anosmia (hereafter called anosmia; median age 56, SD 10.38; 11 women) and 23 where healthy controls (median age 55, SD 7.89; 12 women). There was no significant difference between the groups in age, *t*(41) = 0.52, *p*> .60, *CI* = [-4.17, 7.10]. Only patients with an anosmia duration longer than 6 months, but shorter than 3 years, were included (median duration: 14 months; SD: 9.86; range 7-36 months). Moreover, all patients with anosmia had prior to inclusion in the study been evaluated by an Ear-Nose-Throat (ENT) medical specialist and diagnosed with either idiopathic or post-infectious anosmia. All participants declared themselves as right-handed, without a condition requiring medication, no documented history of neurological diseases, neurodegenerative disorders, or a history of depression, ever experienced head trauma leading to even brief unconsciousness, or had any indication of past trauma based on their in-study acquired T1 image.

Written informed consent was obtained from all participants before inclusion in the study and all aspects of the study were pre-approved by both the ethical review board of the Dresden Medical University (location of data acquisition) and the regional Stockholm ethical review board (analyses location).

#### Olfactory Performance Test

On the day of scanning, all participants (anosmia and control groups) were tested for olfactory performance to assure either functional anosmia or healthy olfactory functions. Individual olfactory performance was assessed using measures of odor detection threshold, cued olfactory identification, and olfactory quality discrimination using the Sniffin’ Sticks testing set (Kobal et al., 1996; Hummel et al., 1997). The Sniffin’ Sticks consist of felt-tipped pens filled with the odorant in question, as described below. Odor detection threshold was assessed for phenylethyl alcohol (PEA), an odorant often used in measures of absolute sensitivity due to its low level of trigeminal irritation (Wysocki et al., 2003), using seven reversals, three-alternative, forced-choice ascending staircase procedure in a 16-step binary dilution series. The arithmetic mean of the four last reversal points was calculated as the individual’s threshold score. For odor quality discrimination, sixteen individual triplets of pens were presented; each consisting of two pens with identical odorants and one with an odorant of different quality (target). Participant’s task was to discriminate the pen containing the odorant different in quality from the other two in a forced-choice task. In the cued odor identification task, olfactory identification performance was assessed with a four-alternative forced-choice cued identification task for sixteen odorants. All three individual tests had a maximum score of 16 and the sum score of the three (TDI, maximum score of 48) was used as a global estimate of olfactory functioning. Three control participants had previously been tested using the same identification test and were therefore not assessed due to the significant recollection of test items; all three had scored within normosmic range and declared as normosmic. As expected, an independent sample Student’s t-test demonstrated that there was a significant difference between the two groups in respect of their TDI olfactory performance scores where individuals with anosmia scored an average of 13.35 (± .50 SEM; range 7-16.25) and controls an average of 35.21 (± .65 SEM; range 29.5-40.5), *t*(39) = 25.36, *p*< .0001, 95 % *CI* = [20.12, 23.60].

### Image acquisition, data processing, and statistical analyses

#### MRI data acquisition parameters

All imaging data were acquired on a Siemens 3T Verio scanner with a 32-channel head coil. A structural T1 weighted scan was acquired (TR 2300, TE 2.89, flip angle 93.8°, field of view 256 x 256, voxel size 1×1×1 mm) together with a 9 minutes long resting state fMRI block consisting of single shot EPI sequences (TR 2300, TE 22ms, flip angle 90°, Field of view 240 x 240, voxel size 2.5 x 2.5 x 2.5 mm) rendering a total of 216 volumes per participant. All participants were instructed to focus their eyes on a marked dot on the scanner boar ceiling, to let their mind wander, and to breathe through their mouth.

#### Multivariate pattern analysis of voxel wise morphology

We assessed whether the gray matter morphology between Anosmia and heathy Controls differ using a multivariate approach. Initially, MRI T1-weighted images were bias corrected and segmented to different tissues, including gray matter, white matter, and cerebrospinal fluid using a unified segmentation approach (Ashburner and Friston, 2005), carried out in the toolbox of Statistical Parametric Mapping software (SPM12). Next, the segmented gray matter (GM) images were iteratively deformed to their mean within the DARTEL framework (Ashburner, 2007). These images were thereafter spatially normalized to MNI space and modulated to preserve the amount GM in the original image. Because we wanted to assess differences of the patterns within regions rather than voxel-based assessments, we used multivariate pattern analysis with a searchlight approach. Specifically, in the first step, GM images were divided into spherical clusters with a radius of 6mm and a linear support vector machine (SVM) was trained for each cluster to generate a whole-brain map of classification accuracy to show how accurately individuals were classified as belonging to the designated Anosmia and Control group (Figure 1A). To produce a generalizable result, we applied a cross-validation with 10 folds which in each fold, the classifier was trained on 38 subjects (half Anosmic and half Control) and tested on the last pair. To avoid biased, the number of individuals in the two groups were equalized. Consequently, for training of a SVM in each fold 19 individuals were semi-randomly selected from a total of 23 controls to match the number of anosmics training samples. At the end of the 10-folds, all individuals (i.e. 20 anosmics and 23 controls) were used at least once to train the SVM. All the training, testing, and k-fold cross validation were implemented in CoSMOS MVPA toolbox (Oosterhof et al., 2016) within Matlab 2018a. An a priori defined statistical threshold for classification accuracy (ACC > 70%; cluster size of k > 100) was applied to detect significant voxels. Post-hoc permutation tests with 5000 resamplings were performed on the clusters surviving the 10-fold cross-validation and p-values calculated as the probability of observed accuracy against permuted accuracy.

**Figure 1.**
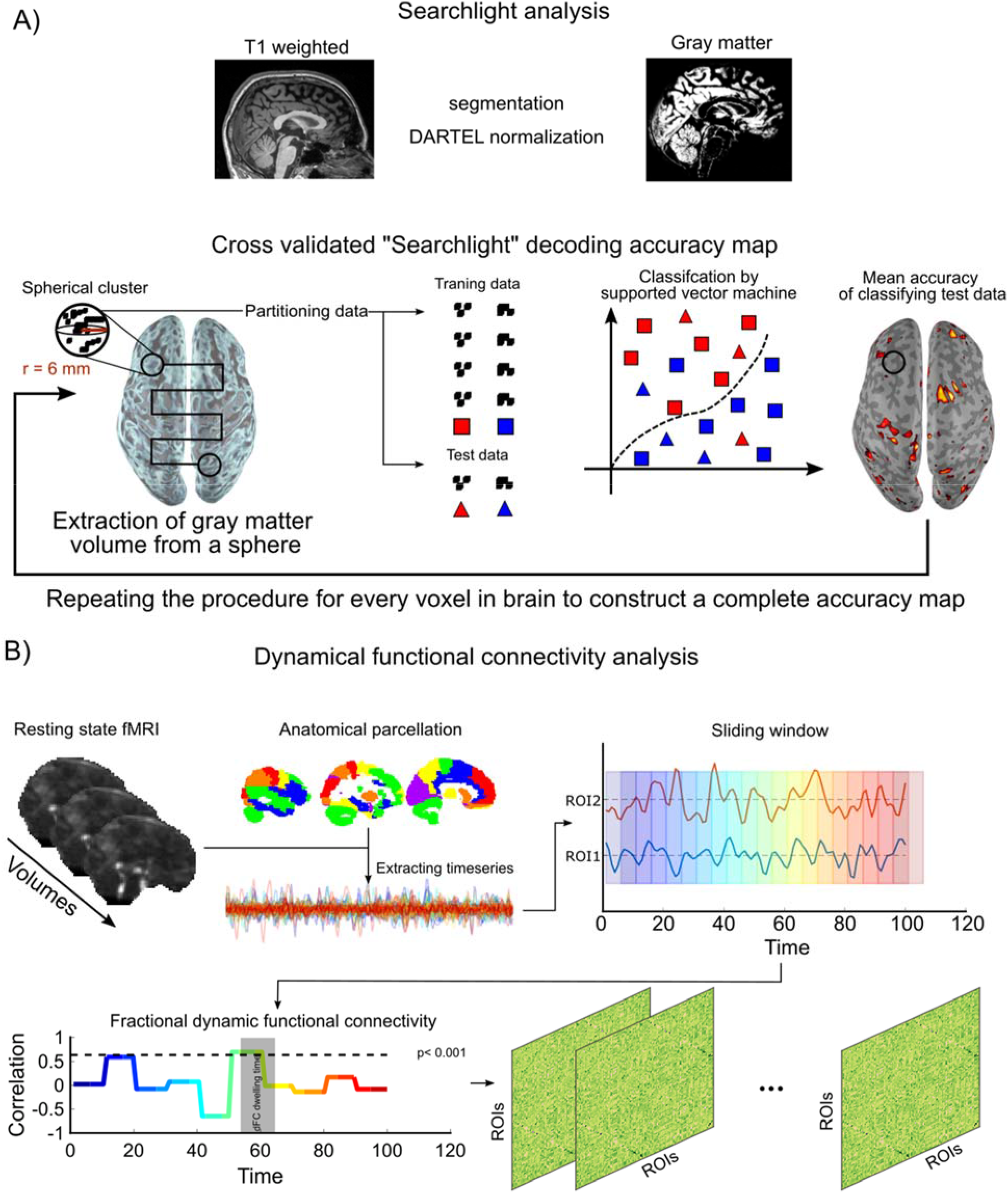
General overview of analysis pipelines. **A)** Anatomical images were pre-processed and voxel-wise gray matter volume values were computed. Group classification accuracy was assessed using a support vector machine with a searchlight approach moving throughout the brain. **B**) The fMRI time series was subdivided into 96 individual ROIs covering the whole-brain. From these ROIs, time series were extracted and temporal correlations from olfactory ROIs to remaining areas were computed within each hemisphere and subsequently averaged between hemispheres. Measure of dynamical functional connectivity was derived from the fraction of the averaged dwelling time to the total scan time.

#### Resting state fMRI

##### Pre-processing

We performed a standard preprocessing pipeline for fMRI data which began with realignment, in which the functional time-series was first realigned to the mean functional image. We then co-registered the structural image to the mean functional image to be able to spatially normalize the functional images, via DARTEL, to MNI-space. The mean functional images delivered the priors for a unified segmentation process and was non-linearly segmented. This yielded the normalization parameters which were applied to all images. All preprocessing steps used the standard SPM12 routines. In addition, beyond using motion parameters as regressors of no interest, we also assessed potential differences between groups in movement throughout the session using frame-wise displacement (FD) (Power et al., 2014). FD is an index that assesses motion of the head between one imaging volume to the next and is estimated as the sum of the absolute values of the differentiated realignment estimate at every time point (Power et al., 2012). A Student’s t-test indicated that there was a nominal, but not significant according to a priory defined statistical threshold, difference between groups in FD values, *t*(41)= 1.92, *p*> .06, 95 % *CI* = [-0.002, 0.085] (mean FD index control: 0.120, SD 0.05; mean FD index anosmia: 0.162, SD 0.09). Note, however, that the confidence interval includes the no difference value.

##### BOLD time Series extraction

First, we created anatomical regions of interest (ROI) to support further analyses. A total of 90 regions of interests (ROI) were extracted from the AAL atlas (Tzourio-Mazoyer et al., 2002) to cover the full brain. In addition, we specifically sought to assess processing within key areas associated with olfactory processing, namely the anterior piriform cortex (APC), posterior piriform cortex (PPC), and the olfactory orbitofrontal cortex (OFC), areas that are poorly defined in the AAL atlas. To this end, we manually created 6 ROIs of the olfactory areas (left and right of APC, PPC, and OFC). The APC and PPC ROIs were based on manual drawings on normalized T1 images from 60 individuals, unrelated to the individuals in this study, using MRICroN (Rorden et al., 2007). Delineation of the piriform cortex was divided into anterior and posterior based on separation between temporal and frontal areas. Mean images of each hemisphere and area were created using the *imcalc* function within SPM 12 and subsequently binarized. For the OFC ROI, we used a functional mask based on a previously published activation likelihood estimate study of neural odor processing; the creation of which is described in detail elsewhere (Seubert et al., 2013). Finally, overlapping voxels were removed from the AAL ROIs using the *imcalc* function (SPM12). This resulted in a total of 96 ROIs. BOLD activation of each ROI was band-passed filtered [.01 - .1 Hz] and averaged across all the voxels within a ROI.

##### Dynamic functional network assessments

Previously, it has been shown that time-resolved resting state brain networks provides a good foundation to further our understanding of dynamic information processing in the brain (Kaboodvand et al., 2020; Zalesky et al., 2014). We sought to determine potential dissimilarities in the functional neural processing at rest between Anosmia and Control using a sliding window paradigm to create inter-regional dynamic functional connectivity (dFC) from the extracted BOLD signal. To remove a potential dependency originating from selected window width, we used a range of windows with different width and averaged the results. We considered windows with width from 5 to 19 with a step of 2 time points (i.e. 8 windows in total) to estimate the dFC for the same ROIs as described above. We also controlled the nuisance effect of movement by regressing out variance that were associated with FD values. Then, dFCs were Fisher Z-transformed to Z-values and to determine significant intervals of connectivity between regions, we applied a threshold on the dFCs where the cutoff value was set based on the critical correlation (*rho_crtical_* = .64; *p*< 0.001); i.e., only connectivity above this value was considered. The fractional connectivity, derived from the threshold dFCs, was calculated for both groups as following: the total interval of dFCs above the threshold was divided by the total length recording within each hemisphere and subsequently averaged across hemispheres for each ROI. Finally, we compared the fractional connectivity across groups from three olfactory regions (APC, PPC, OFC) to the rest of the brain using independent Student’s t-test (Figure 1B).

### Data availability

The data that support the findings of this study are available on request from the corresponding author. The data are not publicly available due to privacy or ethical restrictions because of limitations set in the original ethical permits.

## RESULTS

### Anosmia-associated changes in grey matter volume within olfactory and multisensory processing areas

We first assessed whether we were able to differentiate between individuals with short-term acquired anosmia and healthy control individuals using a SVM with a searchlight paradigm on their extracted whole-brain gray matter volume. Patterns of grey matter volume in multiple clusters throughout the brain could differentiate between anosmia from that of control individuals with a classification accuracy up to 86%. The highest accuracy was found for a cluster within the inferior frontal gyrus (x −47, y 39, z 0) with a classification accuracy of 86%, followed by a cluster within the posterior piriform cortex (PPC) (x 17, y 5, z −20) with a classification accuracy of 81%; Figure 2. In addition, we found that a cluster within the angular gyrus (x −49, y −55, z 48) could significantly dissociate between the two groups with a classification accuracy of 78%; just below the a priori set statistical threshold, there were bilateral clusters within the angular gyrus, Table 1 (see Supplementary Figure S1 for individual results of permutation testing).

**Figure 2.**
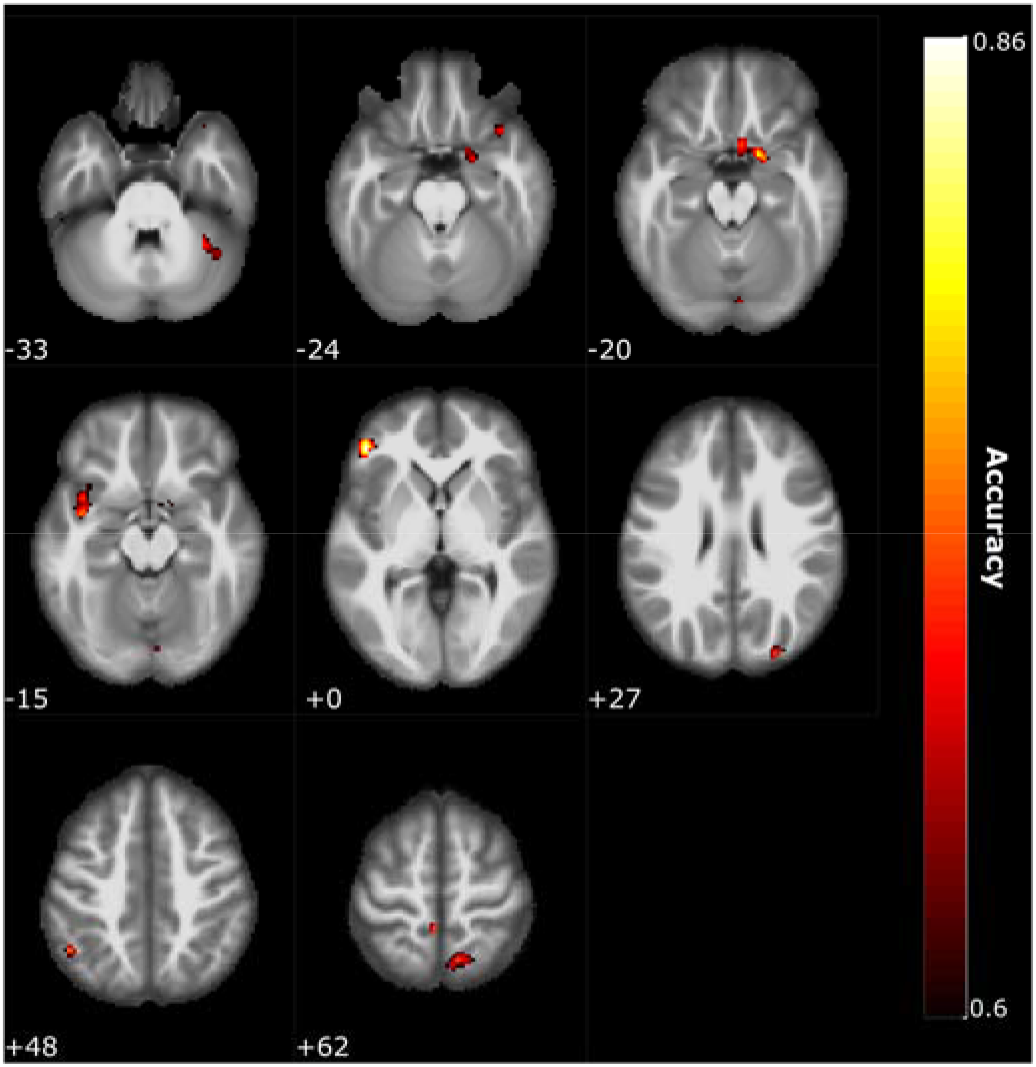
Multivariate pattern analysis. Multiple clusters were found where grey matter volume patterns differentiated between the two groups with classification accuracies between 76 to 86%. Clusters were found within areas associated with olfactory, motor, and multisensory processing (Table 1 and Supplementary Figure 1). Color scheme indicate accuracy level and numbers within figure indicate slice in z-coordinate according to the MNI stereotactic coordinate system.

**Table 1.** List of clusters for 10-fold cross validation SVM with classification accuracy for individuals with anosmia and healthy aged-matched controls based on their gray matter volume patterns. P-values are based on permutation tests with 5000 resamplings.

| Area label | MNI Coordinates |  |  | Cluster Size | Accuracy | P-value |
| --- | --- | --- | --- | --- | --- | --- |
|  | x | y | z |  |  |  |
| Inferior Frontal Gyrus (L) | -47 | 39 | 0 | 266 | .86 | .007 |
| Posterior Piriform Cortex (R) | 16 | 4 | -19 | 334 | .81 | .0046 |
| Precuneus (L) | -4 | -42 | 63 | 113 | .79 | .0168 |
| Superior Occipital Gyrus (R) | 27 | -87 | 27 | 212 | .79 | .0039 |
| Posterior Temporal Pole (L) | -42 | -1 | -15 | 226 | .78 | .0044 |
| Angular Gyrus (L) | -49 | -55 | 48 | 144 | .78 | .0042 |
| Superior Parietal Lobule (R) | 10 | -61 | 61 | 218 | .77 | .0106 |

### Anosmia-induced dynamic connectivity shifts from the olfactory cortex

Having established that short-term acquired anosmia causes differentiable alterations in grey matter volume in, among others, areas associated with olfactory processing, we next assessed whether there were differences in dynamic functional connectivity (dFC). Here, we initially focused on dFC from what is considered core olfactory areas, the anterior- and posterior piriform cortex, as well as the orbitofrontal cortex.

We found that there were significant differences in dFC measured by fractional connectivity between individuals with short-term anosmia and healthy control individuals from the posterior piriform cortex to a range of areas (Figure 3). Among others, participants with anosmia demonstrated stronger dFC from the posterior piriform cortex than control individuals with temporal cortex, *t*(41) = 2.31, *p*< .026, 95% *CI* = [0.004, 0.054] and, given the morphological results reported above, it is of interest to note that individuals with anosmia also demonstrated larger dFC values between posterior piriform cortex and the angular gyrus, *t*(41) = 3.02, *p*< .004, 95% *CI* = [0.006, 0.029]. No significant differences were found from either anterior piriform or orbitofrontal cortex. For the reverse contrast, we did not find any significant differences for the three assessed olfactory ROIs, namely anterior-, posterior piriform cortex, and orbitofrontal cortex (Figure 3).

**Figure 3.**
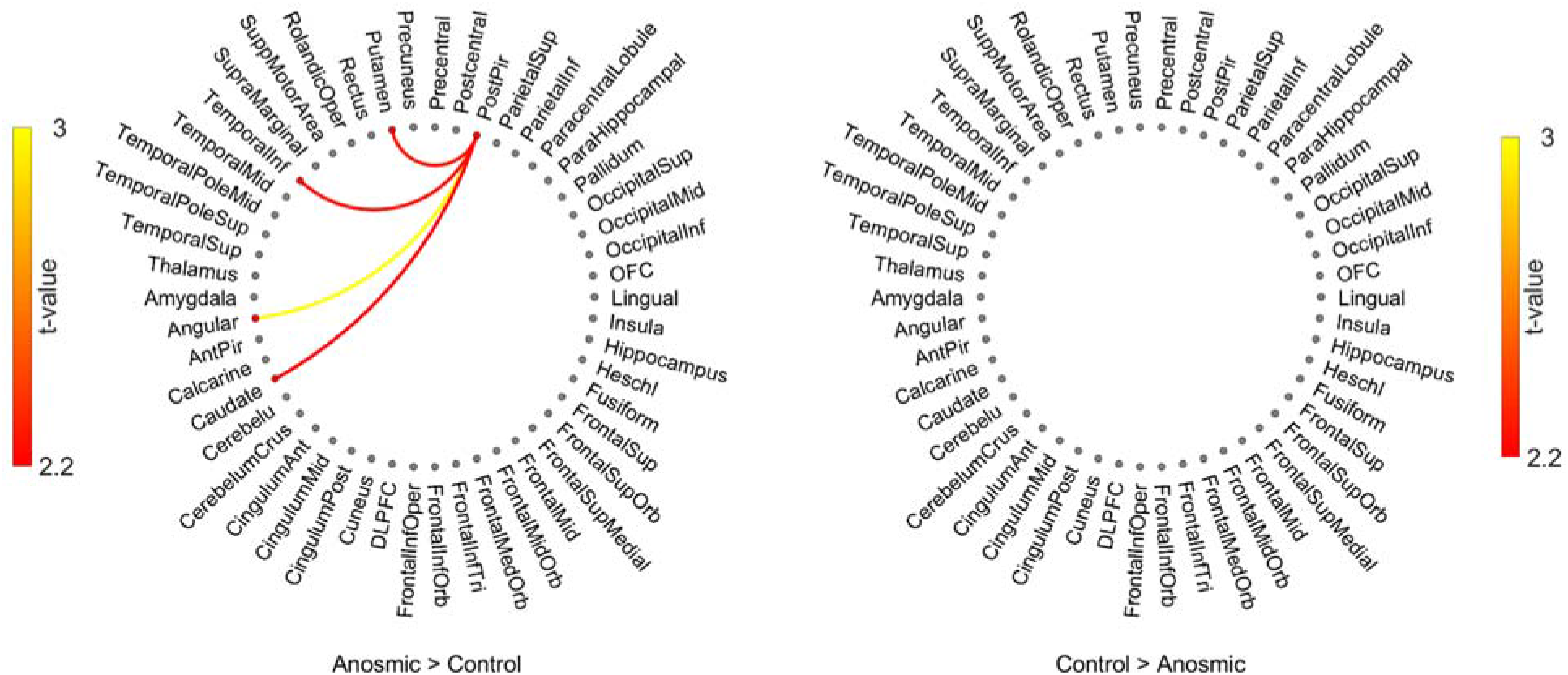
Dynamic functional connectivity from posterior piriform cortex. Dynamic functional connectivity from the posterior piriform cortex to other cerebral regions of interest with significant differences between groups marked with lines. Color of lines represents t-value according to color scale next to each plot. Left side plot shows where individuals with anosmia have stronger inter-regional connectivity and right side plot shows where healthy control individuals have stronger inter-regional connectivity. Only differences with *p*< .05 are shown.

### Anosmia-induced dynamic connectivity shifts from the angular gyrus

Past studies have demonstrated that individuals with anosmia are better at extracting information from multisensory stimuli (Peter et al., 2019) and we recently demonstrated functional gating between posterior piriform cortex and multisensory areas when processing multisensory stimuli (Lundström et al., 2018). Therefore, given the larger GM volume in the angular gyrus on the one hand, and the stronger connection between posterior piriform cortex and angular gyrus on the other, we next assessed whether there was a difference between groups in dFC from the angular gyrus. We found that individuals with anosmia had significantly stronger dFC than healthy controls, among others, between the angular gyrus and somatosensory association cortex in the parietal cortex, the supramarginal cortex, *t*(41) = 2.18, *p*< .035, 95% *CI* = [0.002, 0.049], as well as primary visual area, the Calcarine sulcus, *t*(41) = 2.57, *p*< .014, 95% *CI* = [0.002, 0.020], taste processing areas, the inferior frontal operculum, *t*(41) = 2.06, *p*< .045, 95% *CI* = [0.001, 0.022], as well as the Rolandic operculum, *t*(41) = 2.04, *p*< .047, 95% *CI* = [0.001, 0.025], Figure 4. Finally, in the reverse contrast, we found that healthy controls had only one significantly stronger connections than individuals with anosmia, namely with the inferior temporal cortex, *t*(41) = 3.46, *p*< .001, 95% *CI* = [0.027, 0.101], Figure 4.

**Figure 4.**
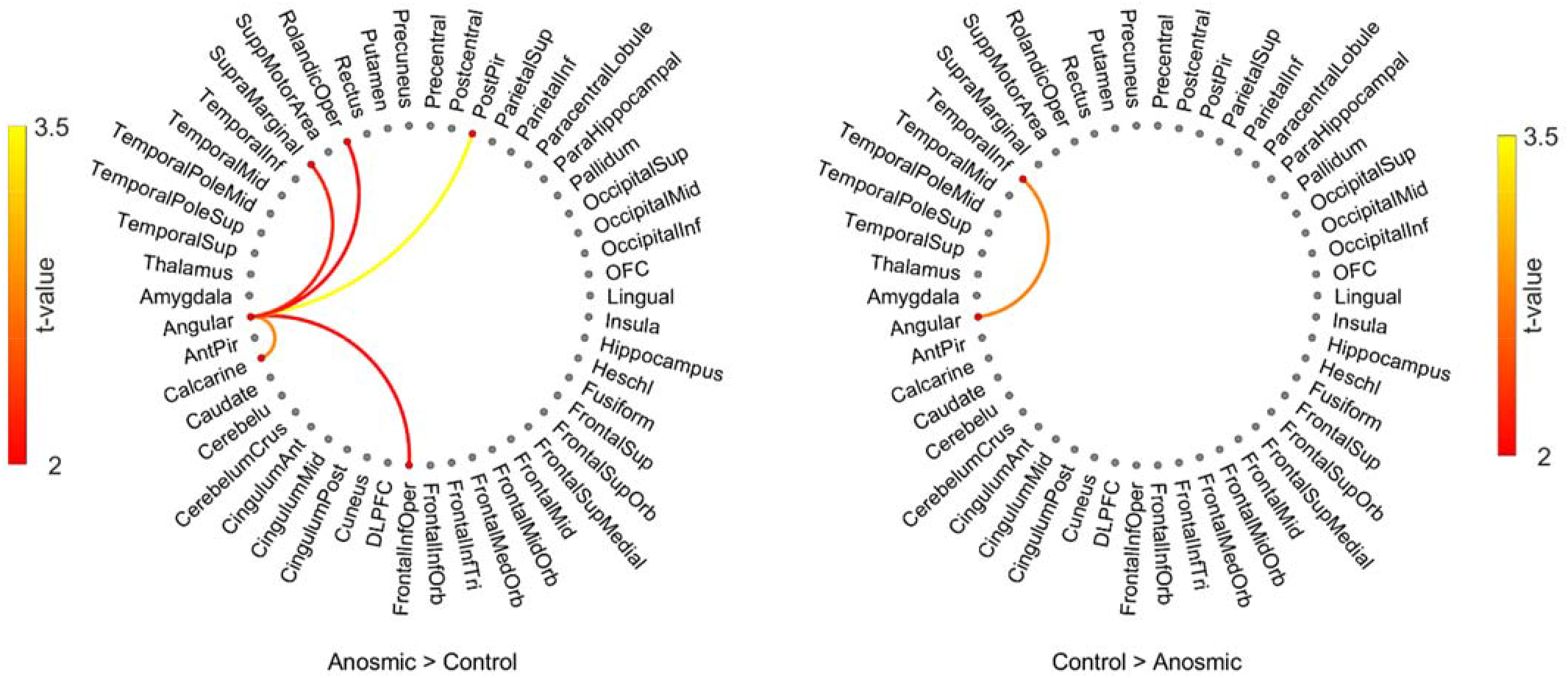
Dynamic functional connectivity from angular gyrus. Dynamic functional connectivity from the angular gyrus to other cerebral regions of interest with significant differences between groups marked with lines. Color of lines represents t-value according to color scale next to each plot. Left side plot shows where individuals with anosmia have stronger inter-regional connectivity and right side plot shows where healthy control individuals have stronger inter-regional connectivity. Only differences with *p*< .05 are shown.

## DISCUSSION

Cerebral plasticity after sensory loss has been well documented in respect of visual and auditory loss but less is known about how the adult brain is affected by the loss of olfactory input. Here we demonstrate that losing the sense of smell in adult age triggers neuroplastic effects that affects both grey matter morphology within, as well as functional connectivity between, cerebral areas. Specifically, we demonstrate that acquired anosmia, i.e. being without olfactory input, during a relatively short time period (average 14 months) is associated with atypical grey matter volume patterns in cortical areas associated with olfactory processing as well as areas beyond the olfactory system. In addition, we demonstrate that acquired anosmia is associated with atypical dFC from the posterior piriform cortex where the individuals with acquired anosmia displayed significantly stronger connection from the posterior piriform cortex to multiple areas within the temporal cortex as well as an area associated with multisensory integration, the angular gyrus; an area where also grey matter volume differed between groups. Finally, we demonstrated that individuals with acquired anosmia had stronger dFC from the angular gyrus to areas primary responsible for basic visual and taste processing.

Compared to the visual system, the olfactory system is dependent on areas that are more widespread as well as more heterogeneous (Mainland et al., 2014; Small, 2004). Sensory deprivation might therefore not have the same effect on olfactory-associated neural areas as areas associated with, for example, visual processing given that the former still receive neural inputs, albeit to a lesser degree, after the sensory loss. Nonetheless, we can here show that several areas demonstrated grey matter volume alterations. Past studies have demonstrated that acquired anosmia change grey matter volume or density in olfactory-related areas, such as the piriform cortex and the orbitofrontal cortex, as well as in non-olfactory-related areas, such as the prefrontal cortex (Bitter et al., 2010; Peng et al., 2013; Yao et al., 2018). Our findings confirm and extend these findings in that we demonstrate a significant difference in an area associated with multisensory integration processing, the angular gyrus. Studies in model species suggests that sensory deprivation alters neuronal populations in multisensory areas (Carriere et al., 2007; Hyvärinen et al., 1981). Moreover, both deaf and blind individuals demonstrate increased recruitment of multisensory areas, such as the angular gyrus and other areas within the posterior parietal cortex, when processing stimuli from intact senses (Bavelier et al., 2001). Along those lines, we recently demonstrated (Peter et al., 2019) that individuals with anosmia are better at integrating multisensory information using an audio-visual integration task that has been linked to the angular gyrus (Zmigrod and Zmigrod, 2015), the angular gyrus is a known integration hub of multisensory stimuli (Regenbogen et al., 2018), and directly linked to the multisensory visual-based integration of olfactory information (Boyle et al., 2007; Gottfried and Dolan, 2003; Regenbogen et al., 2017). One could therefore hypothesize that a lack of olfactory input to these parietal multisensory areas may change the neuronal constellation and promote a more efficient multisensory-based integration of visual and auditory sensory modalities for individuals with anosmia. Measures assessing functional processing of multisensory stimuli were not acquired in the present study but we demonstrate that individuals with anosmia have stronger functional connectivity from the angular gyrus with areas associated with primary visual and taste processing, thus suggesting a functional reorganization that might facilitate improved multisensory processing, much like our past study demonstrated (Peter et al., 2019). Although it is worth noting that the demonstrated enhancement is of a multisensory nature (Peter et al., 2019) and that there is no evidence that anosmia facilitates improved acuity for unisensory visual or auditory stimuli, a clear demonstration of reduced acuity for unisensory taste (Gagnon et al., 2014; Gudziol et al., 2007; Landis et al., 2010) and trigeminal sensations (Frasnelli and Hummel, 2007; Hummel et al., 1996) in individuals with either congenital anosmia or olfactory dysfunction does, however, exist. In light of this, it may be speculated that the increase individuals with acquired anosmia demonstrate in dynamic functional connectivity between the angular gyrus and the operculum, an area linked to both basic taste and trigeminal processing (Albrecht et al., 2010; Veldhuizen et al., 2011), is a so-called hyper-connectivity that in blind individuals is thought to act as a sensory compensatory mechanism (Abboud and Cohen, 2019; Burton et al., 2014; Wang et al., 2014). Increased functional connectivity in response to neurological disruptions may serve as a compensatory mechanism to regulate physiological disturbances, particularly so for hub regions with high connectedness, such as the angular gyrus (Hillary et al., 2014). Indeed, a recent whole-brain modelling study assessing effects of simulated gray matter lesions predicted that, among others, the angular gyrus has a strong hyper-connectivity risk, i.e. would demonstrate an increase in functional connectivity if the whole-brain network faces failures (Kaboodvand et al., 2019). Nonetheless, the direct link between neuroplastic reorganization of posterior parietal cortical areas in acquired anosmia and behavioral improvement in extracting information from multisensory stimuli remains to be confirmed by direct comparisons within a single study. Moreover, it is important to note that results from the support vector machine classification only indicate that there is a difference between groups in gray matter volume patterns, but do not indicate whether either group display more or less grey matter volume.

It is currently not clear what the underlying mechanisms of differences in dFC represent. Whereas static functional connectivity between areas has been demonstrated to be related to high-gamma (~50-150 Hz) inter-regional activity in electrophysiological recordings, dFC seems to reflect connectivity across multiple frequency bands and primarily driven by periods of coordinated high-amplitude coactivation between areas (Karahanoğlu and Van De Ville, 2015; Thompson, 2018). It has further been suggested that an increase in dynamic functional connectivity between areas is represented by an increase in cross-frequency coupling that is responsible for the integration of distributed neural networks (Hutchison et al., 2013). Cross-frequency coupling is strongly linked to perceptual task performance (Fiebelkorn et al., 2013) which is of interest given that dFC has been shown to be a better correlate of behavioral performance than static connectivity (Liégeois et al., 2019). This suggests that differences in dFC between the two populations in the current study could have behavioral implications that should be assessed in future studies combining behavioral and neuroimaging measures. However, whereas the dFC on the individual level was corrected using a conservative critical value and several of the outcomes where supported by analogue morphological findings, the results in the reported dFC group analyses were not corrected for multiple comparisons given the multitude of tests. It is therefore important that these results are replicated before they are extended to future findings.

Individuals with anosmia also demonstrated increased dFC with multiple areas within the temporal cortex. Monosynaptic afferent connections between piriform cortex and the temporal cortex have been demonstrated (Morán et al., 1987) and the inferior temporal cortex and nearby temporal areas have been linked to behavioral performance in tasks tapping object naming and recognition (Tsapkini et al., 2011), odor-visual matching (Olofsson et al., 2013), and odor familiarity (Lundström et al., 2008; Royet et al., 2011; Seo et al., 2013). It is not clear what the potential behavioral implications of this demonstrated stronger dFC connection to multiple areas around in temporal cortex could be for individuals with anosmia; future studies assessing whether acquired anosmia improves behavioral performance in visual object naming and recognition is warranted.

A problem with determining the onset of non-traumatic-dependent olfactory sensory loss is that especially patients with idiopathic olfactory loss have difficulties providing an exact time for when they lost their sense of smell. This means that the anosmia duration stated by the participants should be viewed as an approximation. Nonetheless, on the other extreme of the time-spectra lie individuals who are born without a sense of smell and who can serve as a theoretical contrast to assess potential effects of time. In line with this assumption, and in contradiction with the present results, we recently showed that the piriform cortex in individuals with congenital anosmia did not differ from healthy controls in either gray matter volume, cortical thickness, or functional connectivity to olfactory-associated areas (Peter et al., 2021, 2020; but see Frasnelli et al., 2013). Why there is more evidence of reorganization in primary olfactory regions in individuals with shorter olfactory sensory loss, here with an average of 14 months without olfactory inputs, than in individual with long-term sensory loss is not known. One significant difference between lifelong and acquired anosmia is that the former is characterized by absent or aplastic olfactory bulbs (Abolmaali et al., 2002) whereas the latter demonstrates reduced, yet remaining, olfactory bulbs (Huart et al., 2019). In animal models, ablation of the olfactory bulb has an age-dependent effect on downstream development of the piriform cortex. Ablating the olfactory bulb at birth, hence removing afferent input to the piriform cortex and creating an early olfactory sensory deprivation, has little to no effect on the cortical thickness of the piriform cortex whereas a later removal leads to a significant thinning of the piriform cortex (Friedman and Price, 1986a, 1986b; Westrum and Bakay, 1986). This suggest that differences in the degree of neural plasticity of the piriform cortex between lifelong and acquired anosmia might be linked to the individual’s olfactory bulbs. Whether this hypothesis can be confirmed, and what the potential implications are, need to be studied in model organisms.

In conclusion, our results demonstrate that shortterm acquired anosmia alters grey matter volume in both cortical areas associated with olfactory processing and multisensory integration, as well as increases the dynamic functional connectivity between these areas. Taken together, loss of olfactory input, even at adult age, is associated with changes in morphology of the human brain within, as well shapes the functional connection between, multiple cerebral areas.

## ACKNOWLEDGEMENT

Funding provided by the Knut and Alice Wallenberg Foundation (KAW 2018.0152), awarded to JNL. AA is supported by a grant from the Swedish Research Council (2018-01603).

## Supplementary material

**Figure S1.**
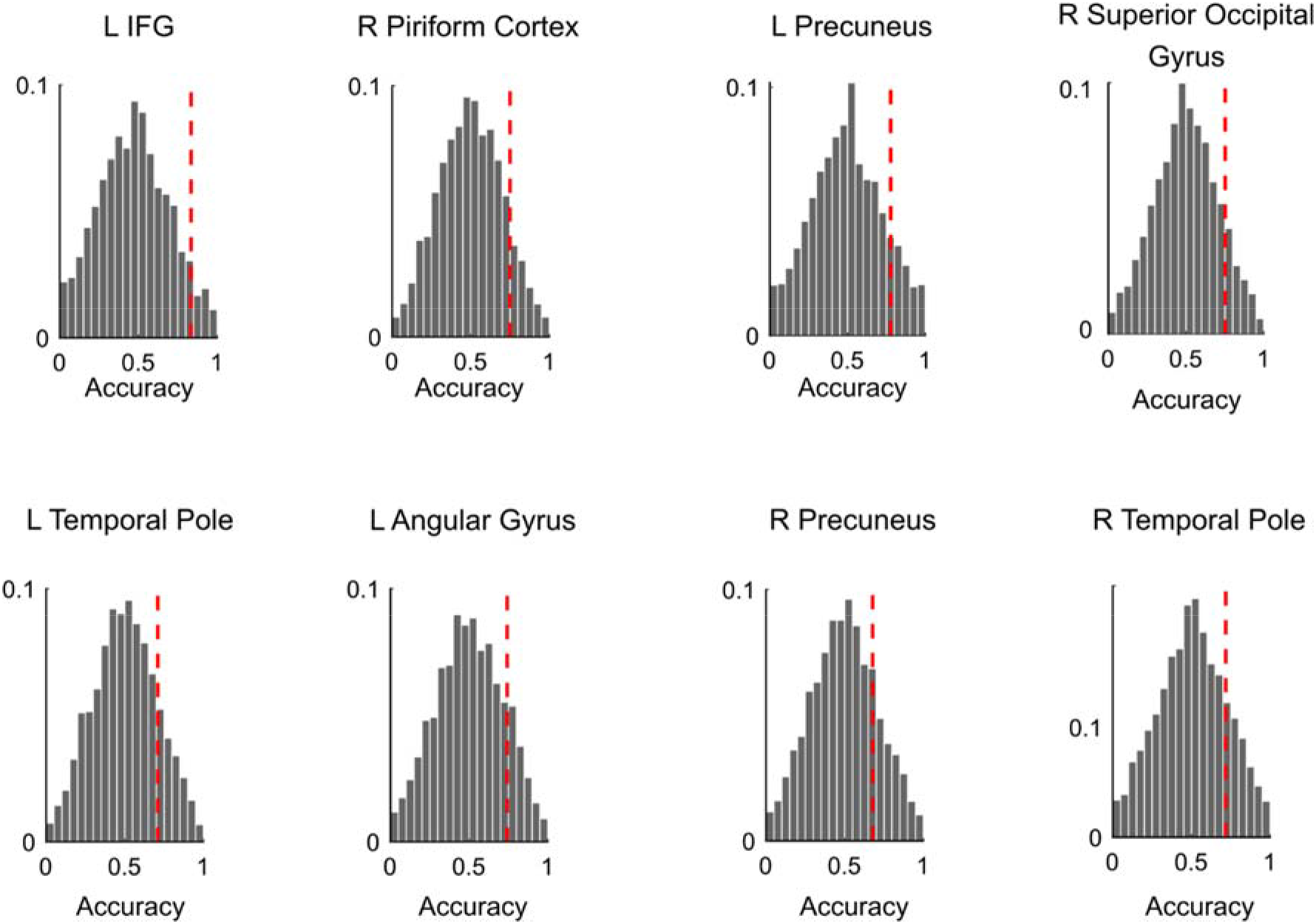
Permutation test of SVM. 5000 permutations performed on the clusters identified by 10-fold cross validation. The distributions show the accuracies of permuted data and red dashed line shows the observed accuracy of the original data.

